# Langerhans Cell-targeted mRNA Delivery: A Strategy for Dose-Sparing and Enhanced Anti-Tumor Immunity

**DOI:** 10.1101/2025.06.25.661517

**Authors:** Klara Klein, Litty Johnson, Ramona Rîca, Mirza Sarcevic, Gabriele Carta, Saskia Seiser, Adelheid Elbe-Bürger, Freyja Langer, Nowras Rahhal, Christoph Rademacher, Robert Wawrzinek, Federica Quattrone, Florian Sparber

**Affiliations:** Cutanos GmbH, Josef-Holaubek-Platz 2 (UZA 2), 1090 Vienna, Austria; Medical University of Vienna, Department of Dermatology, Währinger Gürtel 18-20, 1090 Vienna, Austria; Institute of Science and Technology Austria, Preclinical Facility, Am Campus 1, 3400 Klosterneuburg, Austria; University of Vienna, Department of Pharmaceutical Sciences, Josef-Holaubek-Platz 2, 1090 Vienna, Austria; University of Vienna, Max F. Perutz Labs, Department of Microbiology, Immunology and Genetics, Dr. Bohr-Gasse 9, 1030 Vienna, Austria

**Keywords:** mRNA, lipid nanoparticle, Langerhans cells, Langerin, targeted antigen delivery, C-type lectins, intradermal immunization

## Abstract

Despite the success of mRNA therapeutics, challenges remain in optimizing immune responses and minimizing side effects. Cell-specific antigen delivery may help reduce required doses and improve vaccine efficacy. In this study, we report on the first targeted delivery system for mRNA to a specific subset of skin-resident antigen-presenting cells - Langerhans cells. By functionalizing lipid nanoparticles (LNPs) with a langerin-specific glycomimetic ligand, we achieve selective mRNA delivery to both murine and human primary Langerhans cells with minimal off-target uptake, at the same time resulting in significantly increased mRNA translation. This targeted mRNA delivery not only enhances antigen presentation and T cell responses but also enables dose-sparing and superior anti-tumor immunity compared to conventional immunization in a B16-OVA tumor model. Importantly, our platform’s high compatibility with various lipid nanoparticle formulations offers a flexible and precise tool for skin-directed mRNA delivery.

## Introduction

The SARS-CoV-2 pandemic led to the rapid and widespread adoption of mRNA vaccines, which have saved millions of lives (1). With billions of administered vaccine doses worldwide, both the mRNA itself and its lipid nanoparticle (LNP) carrier have proven to be a safe technology. Since then, mRNA technology is intensely explored for use across various indications, including antiviral vaccines and autoimmune diseases (2). Within this broad landscape, one core focus remains the development of anti-cancer therapeutics - in particular personalized treatment options (3). Antigen-encoding mRNA can be rapidly synthesized and modified to address patients’ specific medical needs while maintaining high quality and purity of vaccine components. But aside from these benefits, significant scope for improvement remains, such as the reduction of side effects and increase of efficacy (4–6).

To fully harness the potential of mRNA technologies, various parameters can be optimized, including mRNA design, LNP composition, and administration route. An effective immune response requires antigen-presenting cells (APCs) to internalize and translate the mRNA. Intramuscular administration of mRNA vaccines leads to an accumulation of mRNA mainly at the injection site and in the liver. However, drainage to lymphoid organs has also been observed and efforts are being made to achieve non-liver mRNA delivery (7–10). To achieve immune cell-specific delivery, LNPs can be modified to target unique surface receptors expressed on APCs. Currently, different types of targeting approaches are explored, including antibodies and their fragments, peptides, oligonucleotide aptamers, or other types of ligands (e.g. proteins and saccharides) (4,11,12). Skin-resident Langerhans cells (LCs) are a subset of dendritic cells (DCs) and considered the “first line of defense” as they are responsible for pathogen recognition, antigen processing, and elicitation of immune responses. This includes activation of T cells, as well as induction of humoral immunity via B and T follicular helper cells (13–17). These features, combined with their location in the epidermis and their ability to mount a systemic immune response, render LCs an attractive access point for immune cell-specific antigen delivery, also with the prospect of minimally invasive administration (18–20).

In this study, we employ a previously described glycomimetic small molecule ligand, mimicking the physiological, multivalent binding interaction with the surface receptor langerin (CD207), expressed by LCs (21). Langerin belongs to the family of C-type lectin receptors (CLR) and interacts with mannose, *N*-acetyl-glucosamine, and fucose residues on the glycocalyx of viruses, bacteria, and fungi, as well as self-antigens (22,23). Upon antigen recognition, langerin shuttles the antigen into endosomal compartments, where a reduced pH causes antigen release, followed by recycling of the receptor back to the cell surface. This process facilitates antigen uptake and presentation, resulting in the elicitation of strong CD8^+^ and CD4^+^ T cell responses. Our langerin-specific ligand has been shown to selectively deliver drugs encapsulated in liposomes or protein antigens to LCs (21,24–26).

Herein, we utilize this langerin-specific ligand, the core component of our “Langerhans Cell-Targeted Delivery System” (LC-TDS), to functionalize LNPs and demonstrate its capacity for cell-specific delivery of antigen-encoding mRNA. Compared to a conventional, non-targeted immunization, we observed superior antigen translation and enhanced *de novo* immune responses *in vivo*. Furthermore, these targeted LNPs allowed for significant mRNA dose-sparing without compromising the anti-tumor response. In conclusion, our findings offer valuable insights towards more efficient and safer mRNA vaccines through immune cell-specific antigen delivery.

## Results

### LC-targeted LNPs enable specific mRNA translation in primary LCs

Given that the LC-TDS preferentially binds to human langerin (21), we used transgenic ‘huLang’ mice, expressing the human CD207 receptor on LCs (27). LC-targeted LNPs (t-LNPs) were prepared using microfluidic mixing by incorporating the LC-TDS-ligand-coupled-PEG2000-lipid (targeting lipid) into four formulations, each containing a different ionizable lipid (SM102, MC3, ALC, or DODMA) (**Figure 1A**). To assess LC-specific LNP uptake and mRNA translation, we incorporated a lipid-bound Alexa Fluor 647 dye (AF647) into the LNP formulations and encapsulated green fluorescent protein (GFP)-encoding mRNA, respectively. Epidermal cell (EC) suspension, containing LCs as the only APC population, were prepared from huLang mice and incubated with t-LNPs (t-SM102, t-MC3, t-ALC, t-DODMA) and their corresponding non-targeted (nt-) controls (**Figure 1B**). All four t-LNP formulations enabled significant LNP uptake (AF647) and mRNA translation (GFP) in LCs, while nt-LNPs did not **Figure 1C, D**). Amongst the tested formulations, t-SM102 was superior in mRNA translation in LCs, followed by t-MC3, t-ALC, and t-DODMA (**Figure 1D**). A similar translation efficacy for t-LNPs was observed even at reduced mRNA concentrations (**Figures S1A, B**). Despite minimal uptake of LNPs by keratinocytes, neither t-nor nt-LNPs induced mRNA translation in these cells (**Figure 1C, Figure S1C**). We further evaluated the targeting specificity by mRNA translation in other DC subsets. Therefore, lymph node (LN) suspension from huLang mice were stimulated with either t- or nt-SM102 LNPs (**Figure 1E**). Upon t-SM102 stimulation, only langerin⁺ LCs, but not langerin⁻ DCs, exhibited dose-dependent mRNA translation. No translation was observed with nt-SM102 (**Figure 1F**).

**Figure 1.**
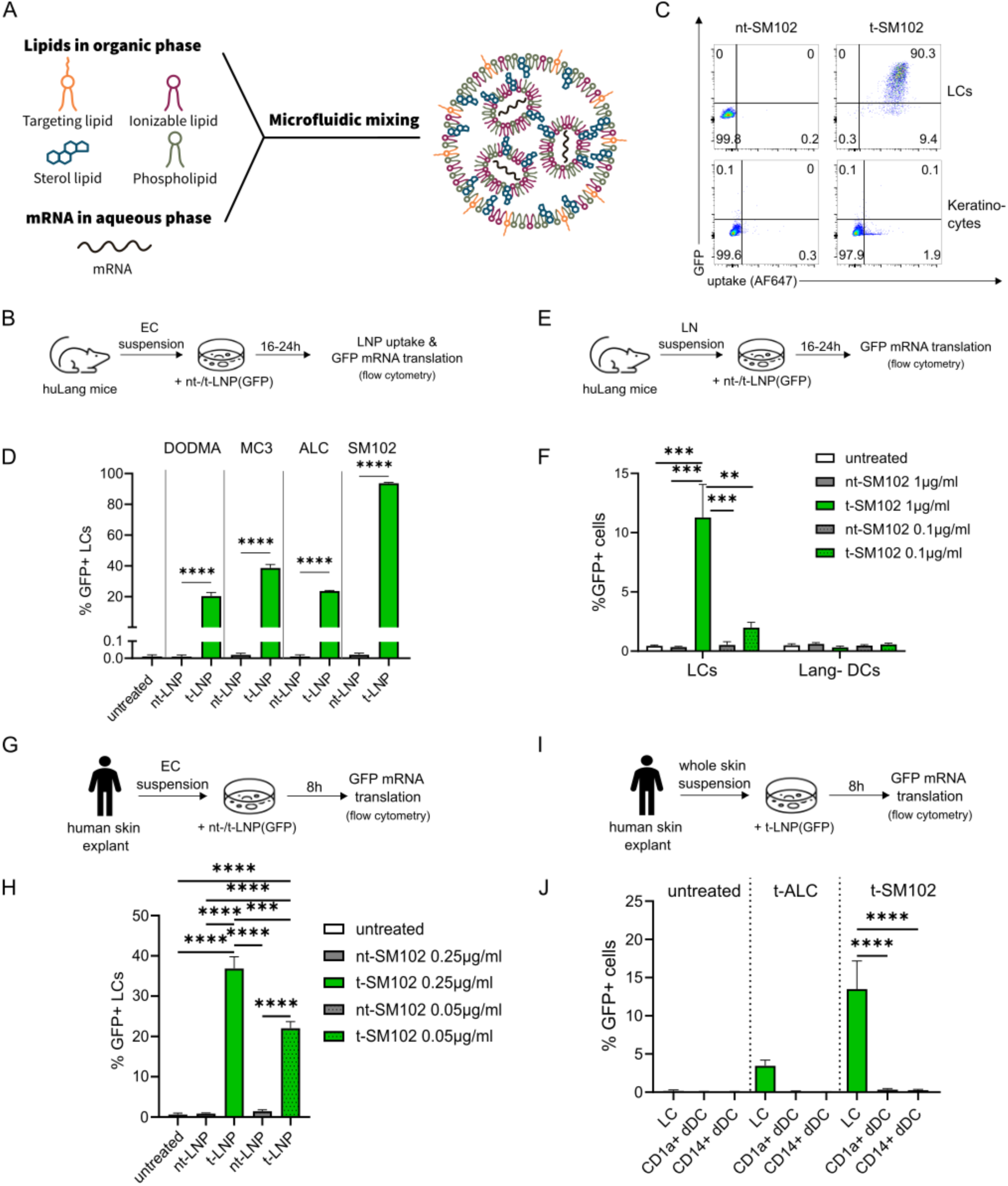
LC-targeted LNPs enable exogenous mRNA translation in primary LCs. (**A**) Representative scheme of the components and structure of t-LNPs used in this study. Particles were generated via microfluidic mixing. An ethanol-based lipid solution comprising of ionizable lipids (Dlin-MC3-DMA, ALC-0315, SM-102 or DODMA), phospholipid (DSPC), cholesterol and LC-TDS-ligand coupled stealth (DSPE-PEG 2000) lipid (targeting lipid) were mixed rapidly with an aqueous phase containing the mRNA in a microfluidic device to formulate t-LNPs. Uncoupled stealth lipid was incorporated instead of the targeting lipid for the generation of nt-LNPs (not depicted). (**B-D**) EC suspension from huLang mice was stimulated overnight with different AF647-labelled nt-/t-LNPs containing GFP mRNA (nt-/t-LNP(GFP)) at a dose of 0.25 µg/ml. LNP uptake and GFP mRNA translation in primary LCs (defined as viable MHC-II^+^ cells) compared to keratinocytes (defined as viable MHC-II^-^ cells) were analyzed by flow cytometry. Data from one representative out of three experiments (n = 3 technical replicates). (**B**) Experimental setup. (**C**) Representative FACS plots for t-/nt-SM102-LNP stimulation, showing uptake (AF647) and mRNA translation (GFP) in MHCII^+^ LCs (upper panel) and MHCII^-^ keratinocytes (lower panel). (**D**) Graph depicts percentages of GFP^+^ LCs upon stimulation with the four different nt-/t-LNPs. (**E-F**) LN suspension from huLang mice was stimulated with nt- or t-SM102-LNPs containing GFP mRNA at a final dose of 1 or 0.1 µg/ml. mRNA translation in murine and human langerin^+^ (mulang^+^hulang^+^) LCs and mulang^-^hulang^-^ DCs (defined as CD45^+^MHC-II^hi^CD11c^hi^ cells) was analyzed by flow cytometry. Data from one representative out of two experiments (n = 3 technical replicates). (**E**) Experimental setup. (**F**) Graph depicts percentages of langerin^+^GFP^+^ LCs and langerin^-^GFP^+^ DCs. (**G-H**) Human EC suspension was stimulated with nt- or t-SM102-LNPs containing GFP mRNA at a final dose of 0.25 or 0.05 µg/ml. Translation of mRNA in CD1a^hi^ LCs (defined as CD45^+^HLA-DR^+^ cells) was analyzed by flow cytometry. Data from one representative out of two experiments (n = 3 technical replicates). (**G**) Experimental setup. (**H**) Graph depicts percentages of GFP^+^ LCs. (**I-J**) Human whole skin suspension was stimulated with t-SM102- and t-ALC-LNPs containing GFP mRNA at a final dose of 0.25 µg/ml (SM102) or 2 µg/ml (ALC). Translation of mRNA in hulang^+^ LCs, hulang^-^CD1a^+^ and hulang^-^ CD1a^-^CD14^+^ dermal DCs (dDCs) (defined as CD45^+^HLA-DR^+^CD11c^+^ cells) was analyzed by flow cytometry. Data pooled from three independent experiments (n = 3). (**I**) Experimental setup. (**J**) Graph depicts percentages of GFP^+^ LCs, GFP^+^CD1a^+^ dDCs and GFP^+^CD14^+^ dDCs. (**D**, **F**, **H**, **J**) Mean ± SEM; one-way ANOVA; ****p<0.0001, ***p<0.001, **p<0.01.

Next, we aimed to validate the functionality and specificity of LC-targeted LNPs *in vivo* using a highly sensitive reporter system. Firefly Luciferase mRNA was encapsulated in t-SM102 LNPs and injected intradermally (i.d.) into the ear pinnae of huLang mice to assess LC-specific translation. Following overnight incubation, LCs and keratinocytes were FACS-sorted from EC suspension, obtained from the injected ears, and luciferase activity was subsequently measured in each cell population (**Figure S1D**). Luciferase expression was detected exclusively in LCs but not in keratinocytes, confirming that intradermal delivery of LC-TDS-functionalized LNPs results in specific mRNA translation in epidermal LCs (**Figure S1E**).

To investigate whether our findings also apply to human LCs, we stimulated human EC suspension (NativeSkin®, Genoskin) with either t- or nt-SM102, containing GFP-encoding mRNA (**Figure 1G**). Again, only t-SM102 induced significant mRNA translation in human LCs (**Figure 1H**), while no translation was detected in keratinocytes (data not shown). To further confirm the targeting specificity in the human setting, we also stimulated human whole skin cell suspension with t-SM102 or t-ALC (**Figure 1I**). Again, both t-LNP formulations yielded GFP mRNA translation exclusively in LCs, with no translation observed in langerin^-^ dermal DCs (CD1a^+^ and CD14^+^ dDCs) (**Figure 1J**).

In summary, our results demonstrate that LC-TDS-functionalized LNPs enable targeted and efficient translation of exogenous mRNA specifically in human and murine primary LCs *in vitro* and upon *in vivo* administration in mouse models. This suggests the potential of the LC-TDS for future applications in skin-targeted mRNA delivery and immunotherapy.

### LC-specific mRNA translation *in vitro* enhances antigen presentation to cognate T cells

To assess whether LC-targeted mRNA delivery translates into functional antigen presentation to T cells, we encapsulated OVA-encoding mRNA using the four aforementioned LNP formulations. Murine EC suspension were co-cultured with naïve CD8^+^ OT-I or CD4^+^ OT-II T cells in the presence of either t- or nt-LNP(OVA) at different mRNA concentrations. T cell proliferation was assessed based on CFSE-dilution (**Figure S2A, D**). All t-LNPs significantly enhanced OT-I T cell proliferation than their non-targeted counterparts, with t-SM102 once again inducing the strongest response across all tested mRNA doses (0.01–1 µg/ml). In comparison, nt-SM102 only triggered robust proliferation at the highest dose (1 µg/ml), while still being inferior compared to its corresponding t-LNP (**Figure S2B-C**).

In OT-II co-culture assays, all t-LNPs (with exception of t-MC3) induced detectable T cell proliferation, though to an overall lesser extent compared to OT-I responses. The corresponding nt-LNPs failed to induce a detectable OT-II T cell proliferation (**Figure S2E-F**). Again, t-SM102 proved to be the most effective formulation, inducing the highest proliferation rate at the intermediate mRNA dose (0.1 µg/ml). Notably, this response decreased at the highest dose, suggesting potential toxicity at elevated mRNA concentrations (**Figure S2F**). In summary, we demonstrated targeted mRNA delivery to LCs to enable effective antigen presentation and significant dose sparing compared to non-targeted delivery, with SM102-containing LNPs showing the highest efficacy.

### LC-targeted mRNA delivery *in vivo* enhances cognate T cell activation

Next, we determined whether enhanced T cell proliferation would also occur following *in vivo* immunization with LC-TDS-functionalized LNPs. Given its previous superiority, we used SM102-containing LNPs and intradermally (i.d.) injected t- or nt-LNPs encapsulating OVA-encoding mRNA (t-/nt-LNP(OVA)) into the ear pinnae of huLang mice. After three hours, EC suspension were prepared from ears and co-cultured with naïve CFSE-labelled OT-I T cells (**Figure 2A**). Administration of t-LNPs significantly enhanced OT-I T cell proliferation (**Figure 2B-C**) and activation (**Figure 2D-E**), compared to nt-LNPs. Notably, robust T cell activation was achieved with as little as 0.25 µg of mRNA that was similar or even stronger than the response seen with 1µg of mRNA.

**Figure 2.**
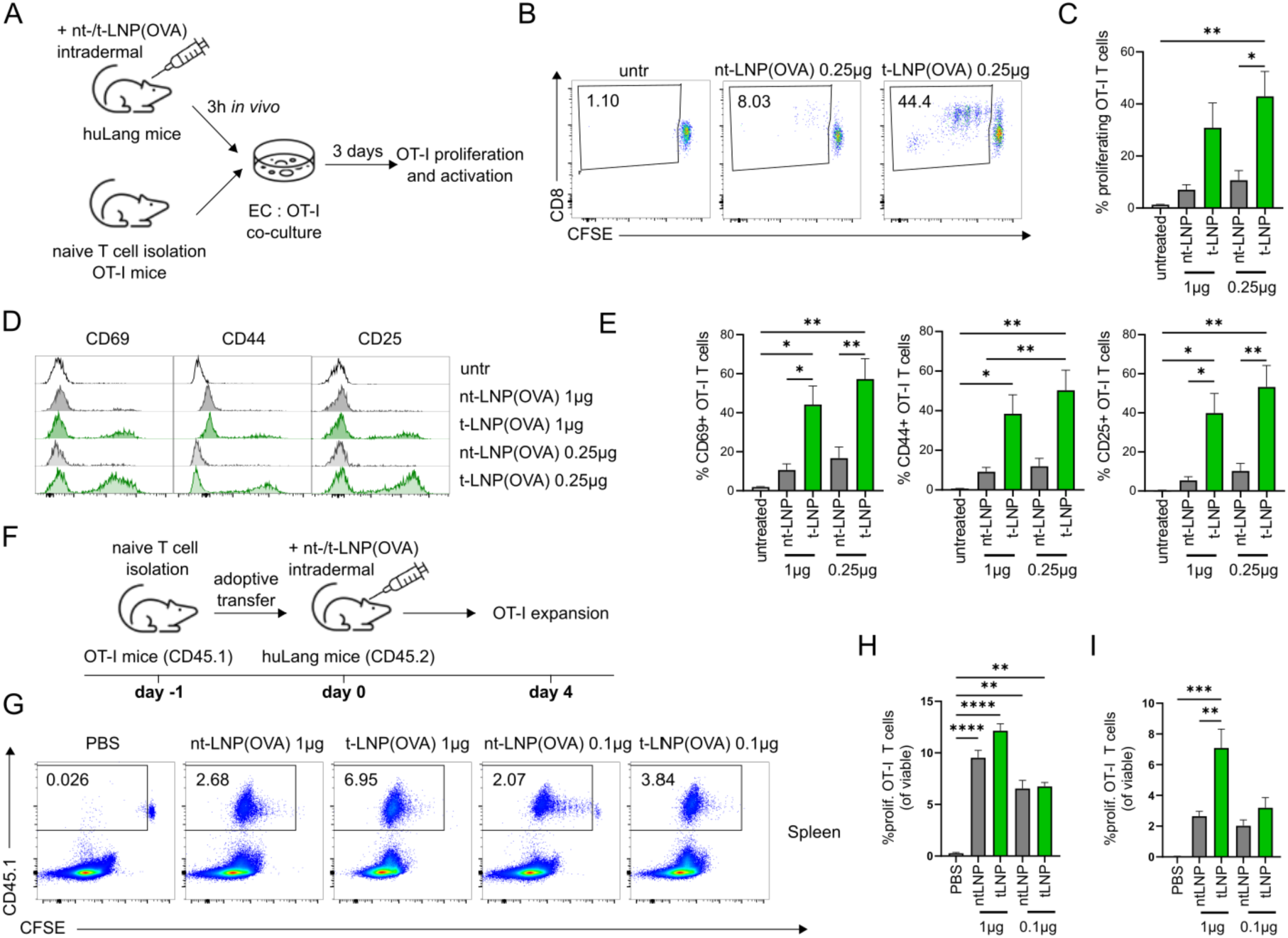
LC-targeted LNPs enhances antigen presentation to OVA specific T cells by primary LCs. (**A-E**) Mice were injected i.d. with nt-/t-LNP containing OVA mRNA (nt-/t-LNP(OVA)) at a final dose of 1 or 0.25 µg. EC suspension was prepared and co-cultured with naïve OT-I T cells. OT-I T cell proliferation and activation was analyzed by flow cytometry. Data from one of two independent experiments (n = 4). (**A**) Experimental setup. (**B**) Representative plots show proliferation of CD8^+^ OT-I T cells (defined as CD45.1^+^TCRVβ5.1/5.2^+^ cells) upon co-culture with EC suspension prepared from mice injected with nt-or t-LNP(OVA) at an mRNA dose of 0.25µg. (**C**) Graph depicts percentages of proliferating CFSE^low^ OT-I T cells. (**D**) Representative histograms show expression of CD69, CD44 and CD25 on CD8^+^ OT-I T cells. (**E**) Graph depicts percentages of CD69^+^, CD44^+^ and CD25^+^ OT-I T cells. (**F-I**) Following adoptive transfer of OT-I T cells, mice were i.d. immunized with nt- or t-LNP(OVA) at a mRNA dose of 0.1 µg or 1 µg. Four days after immunization, OT-I T cell expansion was analyzed in the draining LNs and spleen by flow cytometry. Data from one of two independent experiments (n = 5). (**F**) Experimental setup. (**G**) Representative plots illustrate the gating of proliferating CD45.1^+^CFSE^low^ OT-I T cells in spleen (pre-gated on viable cells). (**H-I**) Graphs depict percentage of proliferating OT-I T cells (defined as CFSE^low^ CD45.1^+^CD3^+^TCRVβ5.1/5.2^+^CD8^+^ cells) out of all viable cells in (**H**) draining LN and (**I**) spleen. (**C**, **E**, **H**, **I**) Mean ± SEM; one-way ANOVA; ****p<0.0001, ***p<0.001, **p<0.01, *p<0.05.

Subsequently, we investigated the antigen-specific T cell response *in vivo*: CD8^+^ OT-I T cells were transferred into huLang mice one day prior to i.d. administration of t- or nt-LNPs. Four days after the immunization, OT-I T cell expansion was analyzed (**Figure 2F**). Surprisingly, transferred CD45.1^+^ OT-I T cells showed strong CFSE dilution in all immunized mice (**Figure 2G**). Accordingly, OT-I T cell expansion was quantified as the percentage of proliferating CFSE^low^ CD45.1⁺ T cells among viable cells. At the dose of 0.1 µg OVA-mRNA, both t- and nt-LNPs induced a comparable CD8⁺ OT-I T cell proliferation. However, t-LNP immunization at a dose of 1 µg OVA-mRNA led to increased proliferation in the draining lymph nodes (dLN) (**Figure 2H**), and more prominently in the spleen (**Figure 2I**), compared to nt-LNP immunization. Overall, these data indicate that LC-specific mRNA delivery upon intradermal immunization results in an enhanced activation of cognate T cells.

### LC-targeted mRNA delivery *in vivo* induces superior *de novo* T cell response

To determine whether the enhanced T cell responses observed upon immunization with LC-TDS-functionalized LNPs extend beyond adoptively transferred OT-I T cells, we investigated the induction of a *de novo* antigen-specific immune response. huLang mice were immunized on day 0 (prime) and 8 (boost) with t- or nt-LNPs, containing OVA mRNA. On day 14 post-prime, T cell responses were analyzing by production of signature cytokines (IFNγ, TNFα and IL-2) following antigen-specific re-stimulation of splenic CD8⁺ and CD4⁺ T cells (**Figure 3A**). Reminiscent of the adoptive T cell transfer setting, immunization with t-LNPs induced a superior CD8^+^ T cell response (**Figure 3B**). This was reflected by higher frequencies of CD8^+^ T cells producing one (**Figure 3C**), two (**Figure 3D**) or three (**Figure 3E**) cytokines. A similar response, albeit to a lesser extent, was also observed for antigen-specific CD4^+^ T cells (**Figure 3F-H**). Taken together, these data demonstrate that LC-targeted mRNA delivery *in vivo* elicits a superior *de novo* T cell response, compared to non-targeted immunization.

**Figure 3.**
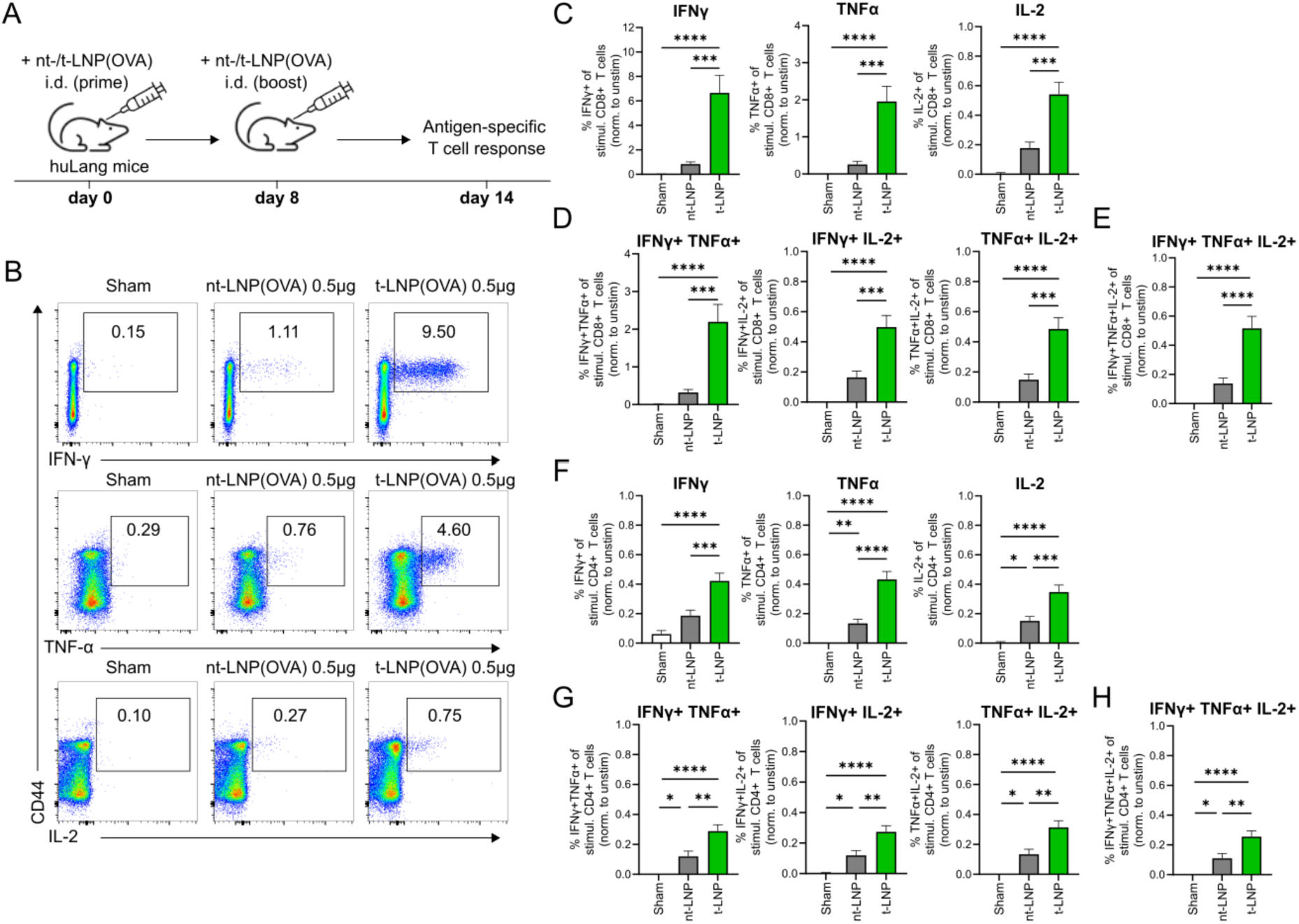
LC-targeted LNPs promote superior endogenous T cell responses. (**A-I**) Mice were immunized i.d. in a prime-boost regimen with nt- or t-LNP(OVA) at an mRNA dose of 0.5µg per immunization. Antigen-specific T cell response was analyzed based on production of signature cytokines by splenic CD8^+^ and CD4^+^ T cells upon antigen-specific restimulation. Data were pooled from two independent experiments (n = 9). (**A**) Experimental setup. (**B**) Representative plots show IFNγ^+^ (top), TNFα^+^ (middle) and IL-2^+^ (bottom) CD3^+^CD8^+^ T cells upon restimulation. (**C-E**) Graphs depict percentages of (**C**) IFNγ, TNFα and IL-2 single, (**D**) double and (**E**) triple cytokine-producing CD3^+^CD8^+^ T cells upon restimulation, normalized to corresponding unstimulated samples. (**F-H**) Graphs depict percentages of (**G**) IFNγ, TNFα and IL-2 single, (**H**) double and (**I**) triple cytokine-producing CD3^+^CD4^+^ T cells upon restimulation, normalized to corresponding unstimulated samples. (**C-E**, **F-H**) Mean ± SEM, one-way ANOVA; ****p<0.0001, ***p<0.001, **p<0.01, *p<0.05.

### LC-targeted mRNA promotes superior anti-tumor immunity with substantial antigen dose sparing

The superior cellular immune response induced by LC-targeted LNPs prompted us to investigate its potential to enhance anti-tumor immunity. To assess this, huLang mice were subcutaneously transplanted with the OVA-expressing B16 melanoma cell line (B16-OVA), followed by three therapeutic immunizations on days 1, 8, and 15 with either t- or nt-LNPs, containing OVA-encoding mRNA (**Figure 4A**). t-LNPs were administered i.d., while nt-LNPs were delivered either i.d. or intramuscularly (i.m.). The i.m. route was selected to reflect a more clinically relevant mode of administration. At a dose of 0.5 µg mRNA, t-LNPs significantly improved survival in tumor-bearing mice compared to i.d. but not i.m. delivered nt-LNPs (**Figure 4B**).

**Figure 4.**
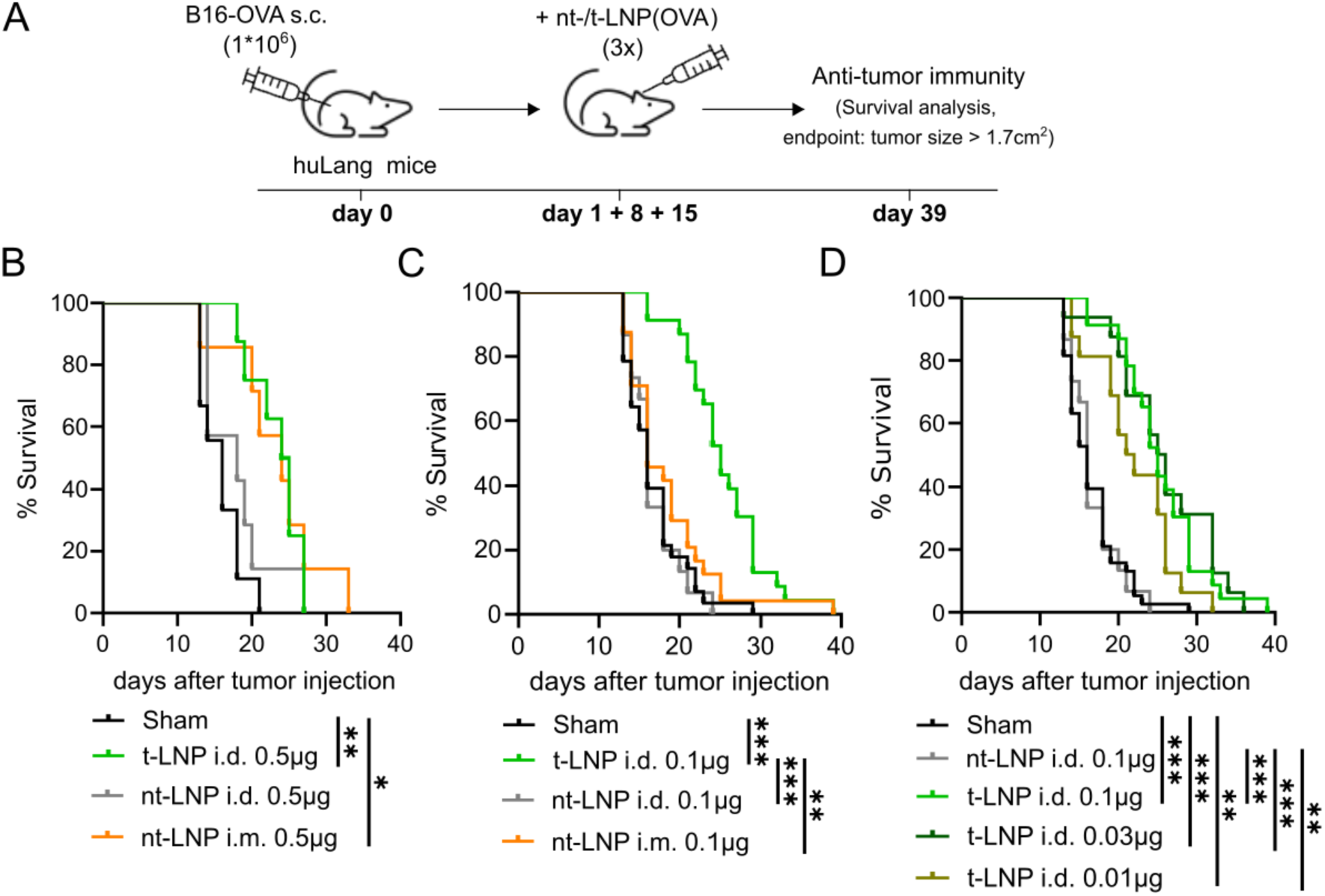
LC-targeted LNPs promote anti-tumor immunity with substantial dose-sparing potential. (**A-D**) Mice were s.c. transplanted with B16-OVA cells followed by three therapeutic immunizations with nt- or t-LNPs, containing OVA mRNA. Survival of animals was analyzed based on tumor growth measurements over time. (**A**) Experimental setup. (**B**) Mice were injected either i.d. with t- or nt-LNPs or i.m. with nt-LNPs at an mRNA dose of 0.5µg per immunization. Survival is depicted as Kaplan Meier plot. Data from one experiment (n = 7-9). (**C**) Mice were injected i.d. with t- or nt-LNPs, or i.m. with nt-LNPs, at an mRNA dose of 0.1µg per immunization. Survival analysis is depicted as Kaplan Meier plot. Data pooled from three independent experiments (n = 15-28). (**D**) Mice were injected i.d. with nt-LNPs at an mRNA dose of 0.1 µg or with t-LNPs at mRNA doses of 0.1, 0.03 or 0.01 µg per immunization. Survival analysis is depicted as Kaplan Meier plot. Data pooled from two to four independent experiments (n = 15-38). (**B-D**) Kaplan Meier plot; log-rank (Mantel-Cox) test (with Bonferroni correction for multiple comparisons), ***p<0.001, **p<0.01, *p<0.05.

To further evaluate the anti-tumor efficacy and dose-sparing potential of LC-targeted mRNA delivery, the mRNA dose was reduced to 0.1 µg per immunization. At this dose, t-LNPs outperformed i.m. administered nt-LNPs, leading to significantly improved survival in tumor-bearing mice (**Figure 4C**). To verify the extent of the antigen dose-sparing, we i.d. administered t-LNPs at an mRNA dose of 0.1 µg, 0.03 µg, and 0.01 µg and compared these to the 0.1µg dose of i.d. applied nt-LNPs. While nt-LNPs failed to elicit an anti-tumor response, t-LNPs induced robust anti-tumor immunity even at the 0.01 µg mRNA dose (**Figure 4D**).

Taken together, this data demonstrates that LC-targeted mRNA delivery induces robust anti-tumor immune responses with a substantial antigen dose-sparing capacity.

## Discussion

Despite the widespread success of mRNA vaccines, particularly regarding the COVID-19 pandemic, ongoing challenges remain in achieving long-lasting immunity whilst minimizing adverse effects. All currently approved mRNA vaccines lack cell-specific delivery and show systemic distribution following intramuscular administration, requiring sufficiently high doses to achieve antigen expression in APCs and subsequent immune activation (4,7,28). These limitations underscore the need for more refined delivery strategies that can direct mRNA expression specifically to professional APCs, thereby improving the quality and magnitude of immune responses while reducing systemic exposure and dose requirements (4).

To address this, we applied the LC-TDS to LNPs and demonstrated its capability for selective mRNA delivery to LCs, a specialized subset of skin-resident APCs. Incorporation of the langerin-specific ligand into LNPs enabled LC-specific translation of the encapsulated mRNA with negligible expression in off-target cells, such as keratinocytes and langerin-negative DCs. This observation was consistent across LNP formulations containing different ionizable lipids, underscoring the robustness and compatibility of our targeting approach. Notably, LNPs incorporating the ionizable lipid SM-102 supported the most efficient mRNA translation in LCs, consistent with previous findings in other cell types (29–32).

The anatomical location of LCs in the epidermis makes them an attractive target for immunization via the i.d. route, leveraging the skin’s high density of APCs. Therefore, this administration route is thought to offer the dual advantages of dose sparing and enhanced immune priming (33–37). The extent of the dose-sparing effect awaits further clinical investigation, as initial studies have shown that fractional i.d. vaccination induces lower immunogenicity compared to standard i.m. administration. Nevertheless, the responses elicited were at levels associated with protective efficacy (38,39).

Our findings demonstrate that i.d. applied LC-targeted LNPs induce potent antigen-specific T cell responses, resulting in robust anti-tumor activity and prolonged survival of tumor-bearing mice, even at remarkably low mRNA doses. We propose that precise targeting can compensate for lower doses and that the LC-TDS has the potential to enhance the dose-sparing capacity promised by i.d. application. This ability is crucial for scalable vaccine deployment and reduction of side effects (36,39).

While being ineffective in eliciting a robust anti-tumor effect at an mRNA dose of 0.5 µg, nt-LNPs administered i.d. still promoted the expansion of adoptively transferred OT-I T cells and initiated a *de novo* antigen-specific T cell response, albeit at reduced levels compared to t-LNPs. This response may be triggered by the uptake of nt-LNPs by APCs other than LCs. Consistently, we cannot entirely exclude that also t-LNPs may interact with other immune cells, yet the use of minute mRNA doses in our tumor model, combined with evidence demonstrating predominant targeting of LCs, suggests that active LC targeting is the primary mode of action underlying therapeutic efficacy.

Administrating vaccines i.d. using conventional needles, as explored in this study, poses practical challenges in clinical settings, including the requirement for proficient personnel and precise injection techniques (36). Advances in needle-free technologies—such as microneedle arrays and needle-free injection devices —may offer a viable path toward broader implementation of skin-based vaccines, improving ease of use and patient compliance (36,40–42). Thus, evaluating the compatibility of LC-targeted mRNA vaccines with these innovative transdermal administration methods could enhance their clinical applicability.

We observed that i.m. administration of nt-LNPs at an OVA-mRNA dose of 0.5 µg produced stronger anti-tumor responses than the same dose delivered i.d. This observation contrasts with the anticipated superiority of the i.d. route based on its higher density of APCs (36). However, our findings align with those from another mRNA vaccine platform, where i.m. compared to i.d. administration resulted in stronger T cell responses (43). Although LCs mainly reside in the epidermis, they can also be found in LNs as migratory LCs. Given that i.m. applied LNPs can drain to LNs (8), it would be valuable to explore whether LC-targeted mRNA delivery synergizes with these muscle-draining pathways. Future studies may uncover new opportunities to leverage the LC-TDS for targeting migratory LCs and to harness potential dose-sparing benefits through the clinically established i.m. route.

In summary, our findings highlight LC-TDS-functionalized LNPs as a versatile and potent platform for targeted mRNA delivery, capable of driving robust T cell-mediated immunity through skin-based immunization at low vaccine doses. This approach holds particular promise for cancer immunotherapy, where T cell responses are key. It also shows potential for prophylactic vaccines, which require both strong humoral as well as cellular responses. As mRNA technologies continue to evolve, the ability to direct antigen expression with cellular and anatomical precision will be essential in optimizing immunogenicity, safety, and scalability. Considering their dose-sparing potential, LC-targeted mRNA vaccines may play a pivotal role in advancing safe and effective future immunotherapies and contributing to global pandemic preparedness.

## Material and Methods

### Human samples and consent

Adult skin was obtained from healthy volunteers after abdominal cosmetic surgery. The study was approved by the local ethics committee of the Medical University of Vienna (ECS 1969/2021) and conducted in accordance with the Declaration of Helsinki Principles. Participants gave their written informed consent. Tissue samples were disinfected with Kodan disinfectant (Schülke & Mayr) and cleaned with phosphate-buffered saline (PBS; Gibco) before cell isolation.

### Mice

Transgenic mice expressing the human langerin receptor (MGI: J:164731) were imported and used under the license agreement from University of Minnesota (27). OT-I transgenic mice (MGI: J:87876) ((44), OT-II transgenic mice (MGI: J:87876) ((45)) and CD45.1 congenic mice (MGI: J:8080) (46) were purchased from The Jackson Laboratory. OT-I or OT-II mice were crossed to CD45.1 mice to generate congenic OT-I/CD45.1 and OT-II/CD45.1 mice respectively. Experimental mice were 8-12 weeks of age and were maintained in the Preclinical Facility (PCF) at the Institute of Science and Technology Austria (ISTA). Animal experiments were approved by the Austrian Federal Ministry for Education, Science and Research (animal protocol number: BMBWF 2022-0.344.288, 2023-0.896.939, 2023-0.930.282, 2024-0-411.154). Animal husbandry and experiments were performed in compliance with national laws and according to the FELASA (Federation of European Laboratory Animal Science Association) guidelines.

### Genotyping

Primer Sequences used for genotyping:

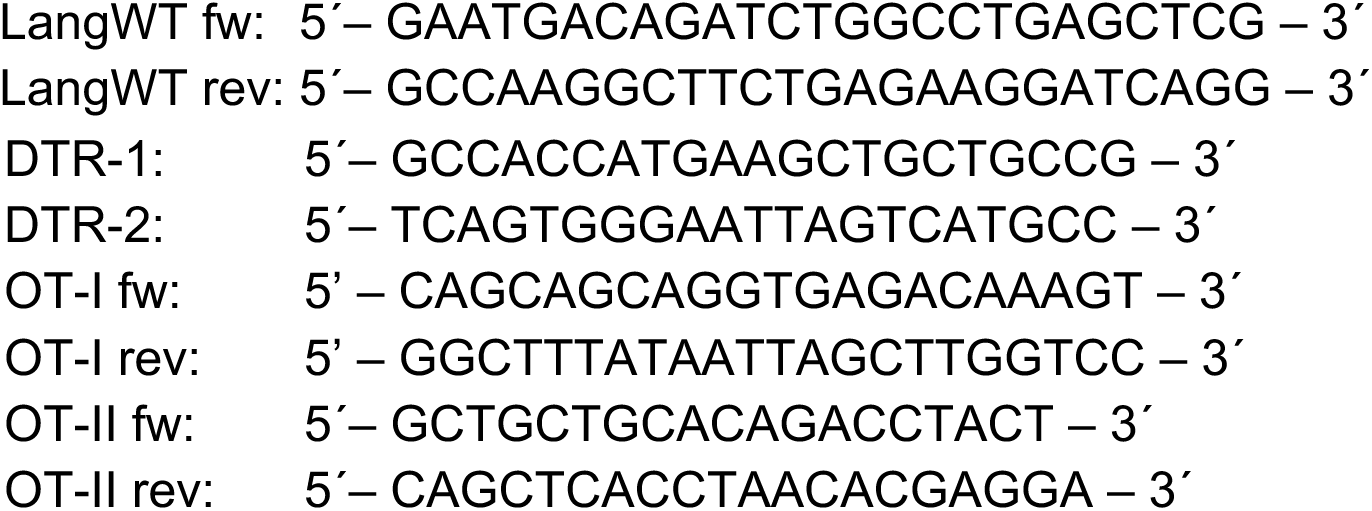

### Formulation and characterization of LNPs

LNPs were formulated using the following ionizable lipids: Dlin-MC3-DMA, ALC-0315, SM-102 (Cayman Europe) and DODMA (NOF Europe). The formulations also included cholesterol (Sigma-Aldrich), DSPC (NOF Europe), DSPE-PEG 2000 (NOF Europe) and the LC-TDS ligand-coupled lipid (targeting lipid). For visualizing cell specific delivery, some LNP formulations contained Alexa Fluor 647 fluorescently labeled lipid conjugate. The mRNA used in these experiments included CleanCap^®^ Enhanced Green Fluorescent Protein mRNA 5moU and CleanCap^®^ Ovalbumin mRNA 5moU (TriLink BioTechnologies) and Firefly Luciferase mRNA (N1-Methylpseudouridine/m1Ψ) (GenScript Biotech Corporation).

The targeting lipid was prepared by conjugating the glycomimetic langerin ligand to DSPE-PEG 2000 NHS (NOF Europe) through an amide reaction. Briefly, 2 mg of the ligand was dissolved in 900 µL 0.1 M sodium bicarbonate buffer at pH 8.4, and the lipid (0.125 molar equivalents) was dissolved in 100 µL dimethylformamid (DMF). The lipid solution was added into the ligand solution, and DMF was subsequently removed by vacuum evaporation (Heidolph). Followed dialysis with a Slide-A-Lyzer cassette (7K MWCO, Thermo Fisher Scientific), the conjugate was then lyophilized to yield the targeting lipid.

The Alexa fluor 647-lipid conjugate was generated by amid-reaction coupling of Alexa Fluor 647 NHS Ester (Thermo Fisher Scientific) to NH2-PEG2000-DSPE (NOF Europe). The lipid was dissolved in DMSO (1 mg per 0.5 mL), and Alexa Fluor 647 (1.5 molar equivalents) was gradually added to the solution. The mixture was allowed to stir overnight at room temperature in the dark. The conjugate was lyophilized using a Alpha 2-4 LDplus, CHRIST and subsequently redissolved in a 0.1 M sodium bicarbonate buffer at pH 8.4. Residual, unreacted dye was removed by dialysis using a Slide-A-Lyzer cassette and the conjugate was lyophilized to yield the fluorescent dye lipid.

For LNP formulation, lipids (Ionizable lipid: DSPC: Cholesterol: targeting-lipid/stealth lipid) were dissolved in 100% ethanol (Sigma-Aldrich) with a molar ratio of 50.5:10.5:37.4:1.6 and mRNA was dissolved in sodium acetate buffer (50 mM, pH 4.5, Invitrogen, Thermo Fisher Scientific). Both solutions were mixed using a microfluidic chip (Wunderlichips GmbH) and the syringe pump 33 DDS system (Harvard Apparatus). The formulated LNPs were then dialyzed overnight using a Slide-A-Lyzer cassette (10K MWCO, Thermo Fisher Scientific) against PBS at pH 7.4 at 4 °C.

After dialysis, size, polydispersity index (PDI) and surface charge of the LNPs were measured by dynamic light scattering (DLS) using the Malvern Zetasizer Pro Malvern Zetasizer Pro. The non-encapsulated and total mRNA content, upon LNP disruption with Triton X-100, were quantified using the Quant-iT^TM^ RiboGreen assay (Thermo Fisher Scientific) following the manufacturers recommendation. The encapsulation efficiency (EE%) was calculated using the formula: Encapsulation efficiency %=RNA (total)-RNA (non-encapsulated)/RNA (total)*100. The physicochemical properties of each formulation used in the paper are reported in the supplementary information (**Table 1-9**).

### Cell culture

B16.F1-OVA (B16-OVA) cells were cultured in Iscove’s minimum essential medium (IMDM) with GlutaMAX (Gibco), supplemented with 5% fetal bovine serum (FBS, Cytiva), 50 μM 2-mercapto-ethanol (Gibco) and 1% Penicillin/Streptomycin (Sigma) at 37°C in 5% CO_2_ (47)

MutuDC 1940-eGFP (MutuDCs) cells were cultured in IMDM with GlutaMAX, supplemented with 10% FBS, 50 μM 2-mercapto-ethanol and 1% Penicillin/Streptomycin at 37°C in 5% CO2 (48).

### *In vivo* immunization strategies

For i.d. or i.m. immunization, mice were anesthetized. by intraperitoneal (i.p.) injection of Ketamine (final: 90 mg/kg) and Xylazine (final: 7.5 mg/kg). The immunization was performed bilaterally either i.d. into the ear pinnae or i.m. into the hind legs (M. gastrocnemius). Different mRNA doses were administered as indicated in the text.

### Generation of murine EC suspension

EC suspension from ear and body skin were prepared as previously described (49). Briefly, ears were cut off split into ventral and dorsal halves. Fur was removed by plucking, and after scraping off the fat tissue, body skin was cut into small pieces. Body skin and dorsal ear halves were incubated epidermal side upwards in 0.6% Trypsin (Sigma) at 37°C. After 20-40min incubation, skin pieces were transferred to FBS (Cytiva) and epidermis was manually detached and further disrupted in complete medium (RPMI1640, Gibco), supplemented with 10% FBS, 1% Penicillin/Streptomycin (Sigma) and L-Glutamine (Sigma). The collected epidermis pieces were incubated for 30min at 37°C in the water bath (shaking) and EC suspension were generated by straining through a 100 µm cell strainer (Sarstedt).

### Preparation of human EC suspension

Human skin explants (NativeSkin access®, Genoskin) were maintained according to manufacturer’s instructions. To prepare EC suspension, the skin was cut in small pieces and incubated epidermal side upwards for 16h at 4°C in complete medium, supplemented with 10 mg/ml Dispase® II (Roche). After incubation, the epidermis was manually detached and further digested with 0.05% trypsin (Sigma) and 0.5 mM EDTA (Invitrogen) in DPBS for 15min at 37°C in 5% CO_2_. The digestion was stopped by adding 0.1% volume of FBS and the tissue was mechanically dissociated. Single cell suspension were prepared by mashing samples through a 100 µm cell strainer.

### Preparation of human whole skin cell suspension

Cell suspension from human whole skin samples were prepared following established protocols(50). Briefly, 4 mm punch biopsies were incubated overnight at 37°C using the human whole skin dissociation kit (Miltenyi Biotec) according to manufacturer’s instructions. Subsequently, mechanical processing was performed using a gentleMACS Octo dissociator (Miltenyi Biotec) with the ‘h_skin_01’ program. Cells were then filtered through a 70 µm cell strainer (Sarstedt).

### Preparation of LN suspension for *in vitro* stimulation

LNs were harvested and collected in Hank’s Balanced Salt Solution (HBSS Ca+ Mg+, Gibco) supplemented with 260 µg/ml of Collagenase D (Roche) and 120 µg/ml of DNase1 (Sigma). After mechanical dissociation, digestion was performed for 25min at 37°C while shaking. Enzymatic digestion was stopped with 10 mM EDTA (Invitrogen) and single cell suspension was prepared by mashing samples through a 100 µm cell strainer.

### *In vitro* stimulation with LNPs

Single cell suspension (0.5-1×10^6^ murine or human ECs, 1.5×10^6^ LN cells or 2.5×10^6^ cells of human whole skin) were seeded in 96 well round-bottom plates (Sarstedt) in complete medium and stimulated with LNPs at a final mRNA concentration of 0.05-2µg/ml for 8-16h at 37°C and 5% CO_2_. Cells were collected and washed before being stained and analyzed by flow cytometry.

### *In vivo* detection of mRNA translation via Luciferase assay

Mice were injected i.d. with t-SM102-LNP containing Luciferase mRNA (GenScript Biotech Corporation) at a dose of 0.25 µg of mRNA/ear. After 16h, EC suspension was prepared from the injected ears and cells were stained for viability and MHC-II expression. MHC-II^+^ LCs and MHC-II^-^ keratinocytes were sorted using a CytoFLEX SRT Cell Sorter (Beckman Coulter). 1×10^4^ sorted cells were seeded into 96-well plates at a final volume of 50 µl in 3-5 technical replicates. 50 µl of ONE-Glo™ (Promega GmbH) were added to each well prior to luminescence signal acquisition with Infinite Pro200 reader (Tecan Trading AG).

### OT-I/II T cell isolation

Naïve CD8^+^ OT-I T cells or CD4^+^ OT-II T cells were isolated using the *naïve* CD8^+^ or CD4^+^ T Cell Isolation Kit (Miltenyi Biotec). Briefly, LNs and spleen were isolated from OT-I/CD45.1 or OT-II/CD45.1 mice and a single cell suspension was generated. After erythrocyte lysis, T cells were enriched using the negative selection kit and labelled with Carboxyfluorescein succinimidyl ester (CFSE, 1:2000, BioLegend) for 20min at 37° C. After washing, cells were resuspended in complete medium for *in vitro* cultures or in DPBS for adoptive transfer experiments.

### EC:OT I/II T cell co-culture

The EC:OT-I/II T cell co-culture was conducted at a ratio of 20:1, respectively, in a total volume of 200 µl/well of complete medium in a 96 well plate (Sarstedt). For co-cultures stimulated *in vitro* with LNPs, 0.5×10^6^ ECs generated from naïve mice were seeded, followed by adding t- and nt-LNP formulations. Subsequently 2.5×10^4^ CFSE-labeled OT-I or OT-II T cells were added to each well. EC:OT-I T cell co-cultures after *in vivo* LNP application were prepared in a similar manner, using EC suspension that was generated from the ears of mice three hours after i.d. LNP injection. The co-culture was incubated for 72h at 37°C and 5% CO_2_. After washing once with DPBS, cells were further processed for flow cytometric staining.

### Adoptive T cell transfer experiments

1×10^6^ CFSE-labelled CD45.1^+^ OT-I T cells were injected i.v. into CD45.2^+^ huLang mice in 100 µl PBS. One day later, mice were i.d. immunized with t- or nt-LNPs at doses of 0.1 or 1 µg of mRNA/mouse. Four days later, OT-I T cell proliferation was analyzed in LNs and spleens.

### Organ preparation for *ex vivo* analysis of T cell response

For *ex vivo* analysis of lymphocytes, draining LNs (auricular lymph nodes) and spleens were harvested and collected. Single cell suspension was generated by mashing organs through a 100 µm cell strainer (Sarstedt). For erythrocyte lysis of spleen samples, cell pellet was resuspended in 1 ml of Red Blood Cell Lysis Buffer (Roche) and incubated for 3 min at room temperature, subsequently stopping the lysis with 5 ml DPBS. After washing, cells were used for flow cytometric staining.

### Endogenous T cell response and *ex vivo* restimulation

Mice were i.d. immunized on day 0 and 8 with t- or nt-LNPs containing OVA mRNA at a dose of 0.5 µg mRNA per immunization. Sham-immunized mice received DPBS. On day 14, antigen-specific T cell responses were analyzed upon *ex vivo* restimulation of splenocytes. MutuDC1940 cells were seeded at a cell concentration of 7.5×10^4^ cells per well the day prior to the restimulation. On the next day, they werepulsed with 50µg/ml OVA protein (Sigma) and 1µg/ml SIINFEKL peptide (Invivogen) 4-6h prior to incubation with 1×10^6^ splenocytes prepared from immunized mice. Incubation with unpulsed MutuDCs served as a control. After 5h, with Brefeldin A (Biolegend) being added for the last 4h, cytokine production by T cells was analyzed by flow cytometric.

### B16-OVA tumor experiments

HuLang mice were s.c. implanted with 0.3×10^6^ B16-OVA cells (in 100µl of PBS) into the right flank. Therapeutic immunizations were performed on day 1, 8 and 15 with nt-LNPs (applied i.d. or i.m. at an OVA-mRNA dose of 0.1 or 0.5 µg per immunization) or t-LNPs (applied i.d. at an OVA-mRNA dose of 0.01, 0.03, 0.1 or 0.5 µg per immunization). Starting with day 5 after tumor transplantation, tumor growth was monitored by caliper measurement every 1-3 days. Survival of tumor bearing mice was monitored, defining a tumor size of > 1.7cm^2^ as the humane endpoint.

### Flow cytometry

Cells were incubated with LIVE/DEAD™ Fixable Near IR or Fixable Yellow Dead Cell Stain Kit (Thermo Fisher Scientific) for 20min on ice, washed and stained with the surface antibody mix for 20min on ice. For the intracellular staining of langerin or cytokines, cells were fixed and permeabilized prior to staining using the BD Cytofix/Cytoperm Kit (BD Biosciences). Analysis was performed using a CytoFLEX S Flow Cytometer (Beckman Coulter) and FlowJo v10.6.1 software (BD Bioscience).

Following antibodies were used for this study: anti-human CD207 (langerin) (MB22-9F5), anti-mouse CD207 (langerin) REAfinity™ (REA822) were purchased from Miltenyi Biotec; anti-mouse I-A/I-E (M5/114.15.2), anti-mouse CD45.2 (104), anti-mouse/human CD11b (M1/70), anti-mouse CD11c (N418), anti-mouse CD103 (QA17A24), anti-human CD45 (HI30), anti-mouse CD3 (17A2), anti-mouse CD62L (MEL-14), anti-mouse CD45.1 (A20), anti-mouse CD25 (PC61), anti-mouse/human CD44 (IM7), anti-mouse CD4 (GK1.5), anti-mouse TCR Vβ5.1, 5.2 (MR9-4), anti-mouse CD8a (53-6.7), anti-mouse CD69 (H1.2F3), anti-human CD1a (HI149), anti-human HLA-DR (L243), anti-human CD11c (Bu15) and anti-human CD14 (HCD14) were purchased from BioLegend.

### Statistics

The applied statistical tests are indicated in the corresponding figure legends. Statistical analysis was performed using Prism 10 (GraphPad). One-way ANOVA followed by Tukey’s multiple-comparison test was used for the comparison of more than 2 groups.

## Author Contribution

Conceptualization: KK, LJ, FQ, FS and RW;

Methodology: KK, LJ, RR, MS, GC, SS, NR, FL, FQ and FS;

Formal Analysis: KK, LJ, RR and FQ;

Writing: Original draft: KK and FS, Review: LJ, RR, AE-B, CR, RW, FQ;

Visualization: KK, LJ, FQ and FS;

Funding Acquisition: RW and FS;

## Acknowledgements

This project was generously supported by “*Seedfinancing”* (P2282679) of the Austrian BMDW and the BMK, handled by the Austrian Wirtschaftsservice (aws), as well as by “*Life Science Call 2022”* (FO999896442) of the Austrian Research Promotion Agency (FFG). We thank Mag. Michael Schunn from the Preclinical Facility (PCF) of the Institute of Science and Technology Austria for his continuous technical support.

---

**Figure S1.**
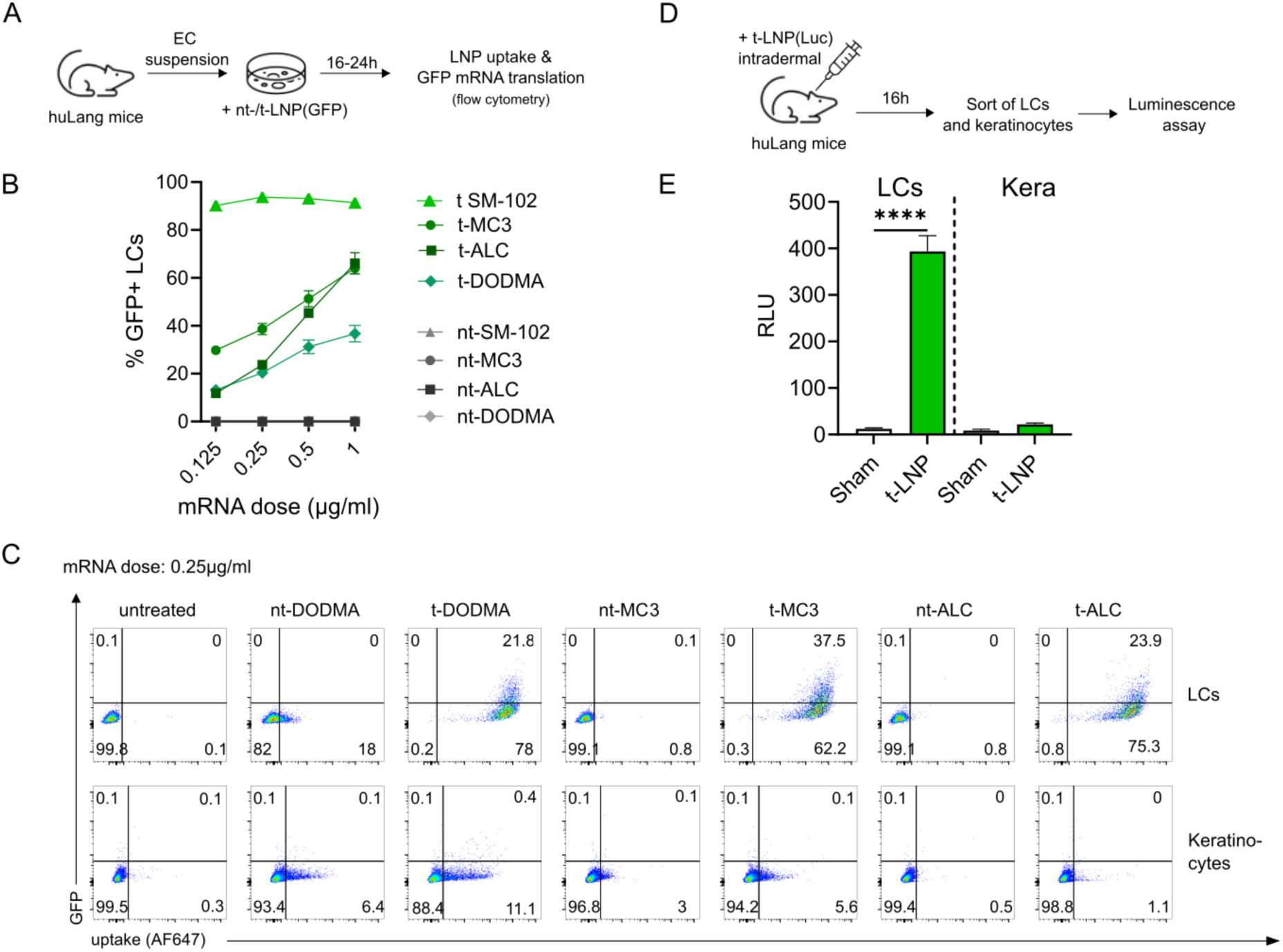
LC-targeted LNPs enable exogenous mRNA translation in primary LCs. (**A-C**) Murine EC suspension was stimulated with nt-/t-LNPs containing GFP mRNA (nt-/t-LNP(GFP)) at final dose of 0.125, 0.25, 0.5 and 1 µg/ml. Translation of mRNA in LCs was analyzed by flow cytometry. Data from one representative experiment (n = 3 technical replicates). (**A**) Experimental setup. (**B**) Graph depicts percentages of MHCII^+^GFP^+^ LCs upon stimulation with different nt-/t-LNPs. (**C**) Representative FACS plots showing uptake (AF647) and mRNA translation (GFP) upon treatment with nt-/t-DODMA, -MC3 or -ALC-LNPs (mRNA dose of 0.25 µg/ml) in MHCII^+^ LCs (upper panel) and MHCII^-^ keratinocytes (lower panel) compared to the untreated control. (**D-E**) Mice were injected i.d. with t-SM102-LNPs containing Luciferase mRNA (t-LNP(Luc)) at an mRNA dose of 0.5µg. LCs and keratinocytes were isolated from the epidermis via cell sorting and luciferase activity was measured in each cell population to assess mRNA translation. Data from one experiment (n = 4-5 technical replicates). (**D**) Experimental setup. (**E**) Graph depicts luminescence signals as RLU (Relative Luminescence Units) in MHCII^+^ LCs (left) and MHCII^-^ keratinocytes (Kera) (right) isolated from immunized and control (sham) mice. Mean ± SEM; unpaired t-test; ****p<0.0001.

**Figure S2.**
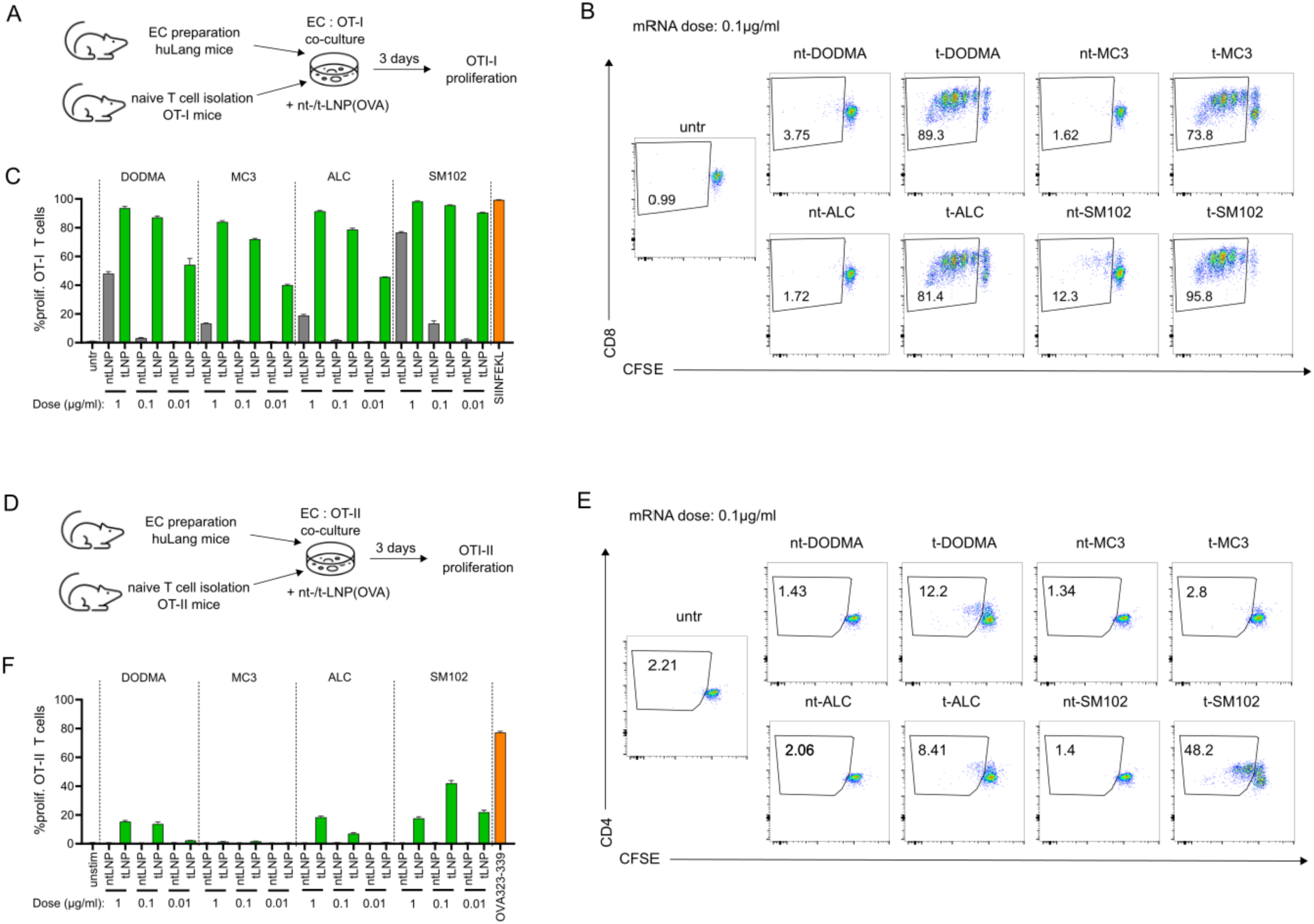
LC-targeted mRNA enhances antigen presentation to OVA specific T cells by primary LCs. (**A-C**) Murine EC suspension was co-cultured with naïve OT-I T cells in the presence of four different nt/t-LNP(OVA) formulations at an mRNA dose of 0.01, 0.1 or 1 µg/ml. After co-culture, OT-I T cell proliferation was analyzed by flow cytometry. Data from one representative experiment (n = 3 technical replicates). (**A**) Experimental setup. (**B**) Representative plots show proliferation of CD8^+^ OT-I T cells (defined as viable CD45.1^+^TCRVβ5.1/5.2^+^CD8^+^ cells). (**C**) Graph depicts percentages of proliferating CFSE^low^ OT-I T cells. Mean ± SEM. (**D-F**) Murine EC suspension was co-cultured with naïve OT-II CD4^+^ T cells in the presence of four different nt/t-LNP(OVA) formulations at an mRNA dose of 0.01, 0.1 or 1 µg/ml. After co-culture, OT-II T cell proliferation was analyzed by flow cytometry. Data from one representative experiment (n = 3 technical replicates). (**D**) Experimental setup. (**E**) Representative plots show proliferation of CD4^+^ OT-II T cells (defined as viable CD45.1^+^TCR Vβ5.1/5.2^+^CD4^+^ cells). (**F**) Graph depicts percentages of proliferating CFSE^low^ OT-II T cells. Mean ± SEM.

